# ForestScanner: A mobile application for measuring and mapping trees with LiDAR-equipped iPhone and iPad

**DOI:** 10.1101/2021.12.11.472207

**Authors:** Shinichi Tatsumi, Keiji Yamaguchi, Naoyuki Furuya

## Abstract

1. Ground-based light detection and ranging (LiDAR) is becoming increasingly popular as an alternative means to conventional forest inventory methods. By gauging the distances to multiple points on the surrounding object surfaces, LiDAR acquires 3D point clouds from which tree sizes and spatial distributions can be rapidly estimated. However, the high cost and specialized skills associated with LiDAR technologies have put them out of reach for many potential users.
2. We here introduce ForestScanner, a free, mobile application that allows LiDAR-based forest inventories by means of iPhone or iPad with a built-in LiDAR sensor. ForestScanner does not require any manual analysis of 3D point clouds. As the user scans trees with an iPhone/iPad, ForestScanner estimates the stem diameters and spatial coordinates based on real-time instance segmentation and circle fitting. The users can visualize, check, and share the scanning results *in situ*.
3. By using ForestScanner, we measured the stem diameters and spatial coordinates of 672 trees within a 1 ha plot in 1 h 39 min with an iPhone and in 1 h 38 min with an iPad (diameter ≥5 cm; detection rate = 100%). The diameters measured by ForestScanner and a diameter tape were in good agreement; R^2^=0.963 for iPhone and R^2^=0.961 for iPad. ForestScanner and a conventional surveying system showed almost identical results for tree mapping (assessed by the spatial distances among trees within 0.04 ha subplots); Mantel R^2^=0.999 for both iPhone and iPad. ForestScanner reduced the person-hours required for measuring diameters to 25.7%, mapping trees to 9.3%, and doing both to 6.8% of the person-hours taken using a dimeter tape and the conventional surveying system.
4. Our results indicate that ForestScanner enables cost-, labor-, and time-efficient forest inventories. The application can increase the accessibility to LiDAR for non-experts (e.g., students, citizen scientists) and enhance resource assessments and biodiversity monitoring in forests worldwide.

## 1 INTRODUCTION

Forest inventory has been an essential component of forest management for over 200 years and constitutes the basis for forest ecology and environmental policymaking today (Corona, Chirici, McRoberts, Winter, & Barbati, 2011; Newnham et al., 2015). With increasing recognition of forest multifunctionality, the use of forest inventory data has expanded from quantifying timber productivity to assessing forest biodiversity and carbon sequestration (Corona et al., 2011). Spatial data of trees in forest inventory plots allow detailed analyses of forest dynamics (Tatsumi, Cadotte, & Mori, 2019; Tatsumi, Iritani, & Cadotte, 2021) and species coexistence (Kunstler et al., 2015; Tatsumi, Owari, & Mori, 2016). As such, forest inventories offer a fundamental step toward studying forest ecology and understanding the benefits forests provide to human societies.

Ground-based light detection and ranging (LiDAR), also known as terrestrial, mobile, or personal laser scanning, is becoming increasingly popular as an alternative means to conventional forest inventory methods (Newnham et al., 2015; Liang et al., 2016). By gauging the distances to multiple points on the surrounding object surfaces, LiDAR acquires 3D point clouds from which trees’ sizes and spatial distributions can be rapidly estimated. However, the high cost of LiDAR devices, typically priced at over US $40,000, has put them out of reach for many potential users (Eitel, Vierling, & Magney, 2013). The heaviness of LiDAR devices has also been a challenge, making their transportations to/within some areas difficult as well as adding shipping costs (Eitel et al., 2013). The need for specialized computer programs has further limited the LiDAR user base (Dassot, Constant, & Fournier, 2011; Bunting, Armston, Lucas, & Clewley, 2013). Broader acceptance of LiDAR as a valid alternative to conventional methods thus requires affordable and efficient hardware and software (Newnham et al., 2015).

Since 2020, Apple Inc. (Cupertino, CA, USA) have introduced a LiDAR sensor in some iPhone and iPad models, namely iPhone 12 Pro, iPhone 12 Pro Max, iPhone 13 Pro, iPhone 13 Pro Max, iPad Pro 2020, and iPad Pro 2021 (as of March 2022). Compared to other LiDAR devices in the market, the LiDAR-equipped iPhones/iPads are available at cheaper prices (US ≥$749) and are lighter in weight (187–684 g). These iPhones/iPads have been found suitable for acquiring 3D point clouds in forests (Gollob, Ritter, Kraßnitzer, Tockner, & Nothdurft, 2021; Mokroš et al., 2021). However, to derive tree information (e.g., stem diameter) from the acquired point clouds, one must conduct post-analyses on a separate device using multiple software, due to the current lack of dedicated iPhone/iPad application programs.

We here introduce ForestScanner, a free application that allows LiDAR-based forest inventories by means of iPhones/iPads. As the user scans trees with the device by walking in the field, ForestScanner estimates the stem diameters and spatial coordinates in real-time (Figure 1; see also Video S1 at FigShare, https://doi.org/10.6084/m9.figshare.17161823.v1). The users can visualize, check, and share the scanning results *in situ*. These features of ForestScanner allow rapid forest inventories and increase the accessibility to LiDAR technologies for non-experts.

**Figure 1.**
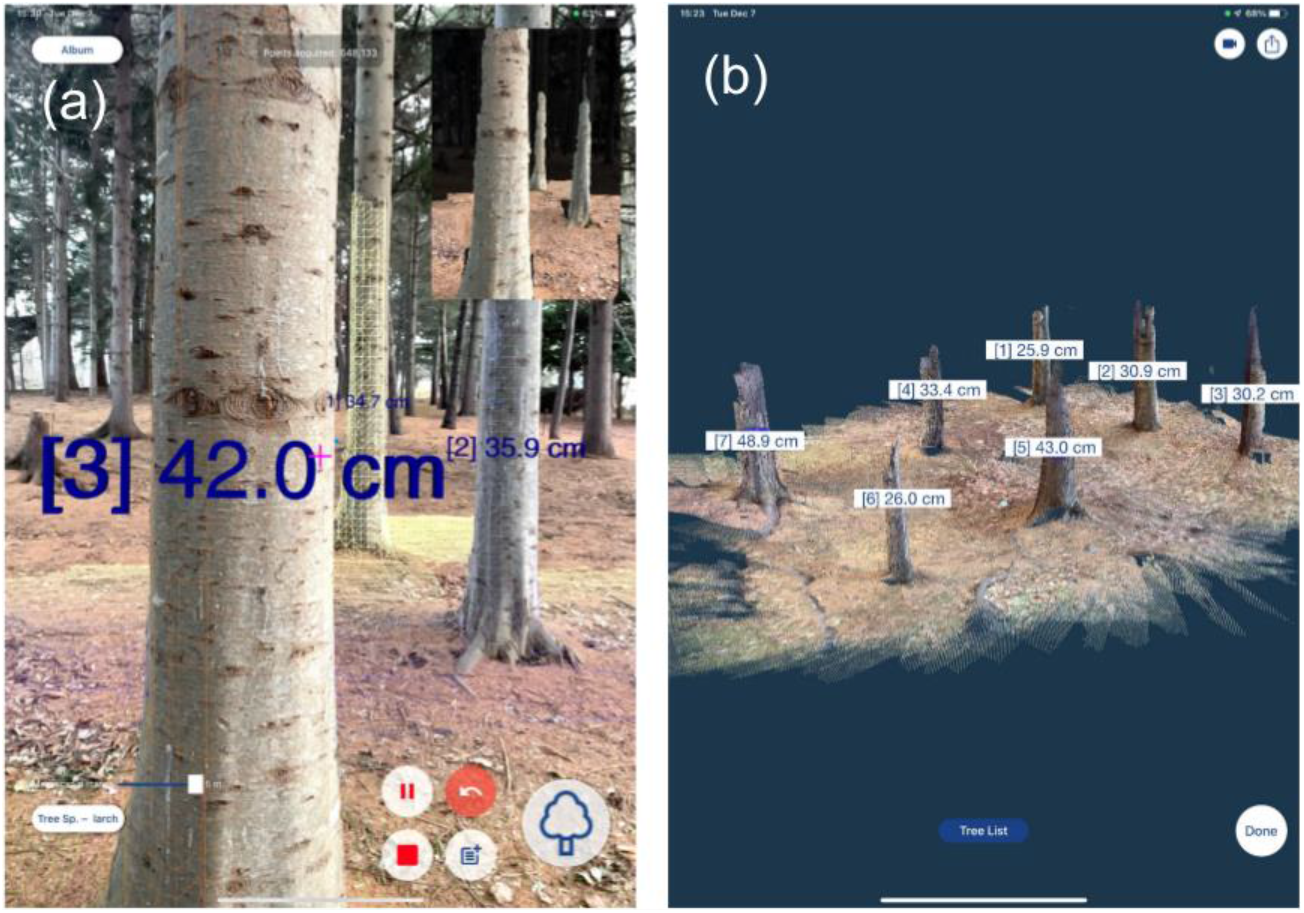
Screenshots of ForestScanner. (a) As the trees are scanned, stem diameters appear instantly on the screen in an augmented-reality manner. (b) An example of point cloud and tree diameters obtained by ForestScanner. See also Video S1 (https://doi.org/10.6084/m9.figshare.17161823.v1) for a demonstration of the app’s usage.

## 2 FORESTSCANNER APP

### 2.1 Description

The ForestScanner application is available for free on the App Store (https://www.apple.com/app-store). The application operates on iPhones/iPads that have a time-of-flight LiDAR sensor. The maximum scanning distance of the sensor is 5 m. As the user moves the device, ForestScanner acquires a 3D point cloud of the surrounding object surfaces. When using ForestScanner, the point cloud and 3D triangle meshes appear on the screen in real-time, allowing users to visually recognize the scanned surfaces (Figure 1a; Video S1). The point cloud is colorized with RGB information collected by the camera on the device. During the scan, ForestScanner keeps track of the device’s relative coordinates from the starting point based on the inertial measurement unit (IMU) (i.e., dead reckoning). Other technologies such as SLAM are not used for tracking the relative coordinates. The absolute location (i.e., geographic coordinates) of the starting point is determined by the GNSS receiver built into the iPhone/iPad.

Similar to other existing methods for LiDAR-based forest measurements (e.g., Dassot et al., 2011; Pueschel et al., 2013; Calders et al., 2015), ForestScanner performs stem detections and circle fitting as separate tasks. First, for a given target tree, real-time instance segmentation is conducted by means of the YOLACT++ fully-convolutional network model (Bolya, Zhou, Xiao, & Lee, 2022). The YOLACT++ model used in ForestScanner has been trained from scratch with a custom dataset consisting of 391 tree images and annotations (Figure S1). The model’s average precision, calculated using multiple thresholds of intersection over union (IoU; 0.50–0.95) (Padilla, Netto, & Da Silva, 2020), was 75.0.

ForestScanner then fits a circle to the cross-section of a given tree by minimizing the sum of squares (*SS*) defined as:

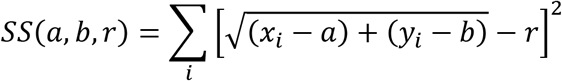

where *a* and *b* are the center coordinates of the circle (i.e., the spatial position of the target tree), *r* is its radius, and *x*_*i*_ and *y*_*i*_ are the coordinates of data point *i*. ForestScanner estimates *a*, *b*, and *r* that minimize *SS* by means of Levenberg-Marquardt (LM) algorithm. The LM is a geometrical algorithm that is commonly used for circle fitting in LiDAR-based forest measurements (e.g., Bu & Wang, 2016).

### 2.2 Usage

- Tap the “Record” button on the screen to start scanning. A crosshair symbol, a slider bar, and the “Tree”, “Pause”, “Back”, “Note”, “Stop”, “Tree species”, and “Album” buttons will appear on the screen (Figure 1a, Video S1).
- While walking in the field, scan target trees individually from a distance of 0.3 m to 5 m. Use the slider bar to fix the maximum scanning distance.
- For each tree, overlay the crosshair symbol at the height where the diameter should be measured (e.g., diameter at breast height; DBH). Then tap the “Tree” button to identify the stem. The stem diameter will appear on the screen in an augmented-reality manner (Figure 1a). The recommended lower threshold of target tree diameter is 5–10 cm (Gollob et al., 2021; Mokroš et al., 2021), as the current iPhone/iPad LiDAR sensor has limited capability of detecting objects that are a few centimeters in size (Vogt, Rips, & Emmelmann, 2021).
- The user can scan multiple trees in a single survey by moving from one tree to the next while scanning. Alternatively, the user can tap the “Pause” button, move to the next tree, and then tap the “Record” button to resume. During the pause, the LiDAR sensor stops scanning while the IMU continues to keep track of the device’s location. Using the pause function helps to reduce the file size of the point cloud.
- Tap the “Tree species” button to enter the tree species (optional). The tree species can be entered by using either the screen keyboard or voice input (i.e., iPhone/iPad’s microphone).
- Tap the “Back” button to delete the tree previously scanned.
- Tap the “Note” button to add a note about the tree previously scanned.
- Tap the “Stop” button to end the survey. The acquired point cloud will be displayed on the screen (Figure 1b).
- After the survey, tap the “Tree List” button to show the list of tree ID, diameters, geographic coordinates, tree species, and notes. The list can be sent to other devices by email or AirDrop.
- Tap the “Album” button to show the results of previous surveys.

## 3 FIELD APPLICATIONS

### 3.1 Data collection and analyses

We applied ForestScanner to a 1 ha (100 m × 100 m) plot composed of twenty-five 0.04 ha (20 m × 20 m) subplots in the Hokkaido Research Center, Forestry and Forest Products Research Institute, Japan (42°59′57” N, 141°23′29” E) (Figure 2a). The plot encompasses mosaics of multiple forest types including conifer and broadleaf plantations of different stand ages and natural secondary forests (Figure 2b). The plot had almost no understory vegetation except in the secondary forests where dwarf bamboos (height ≤1.3 m) covered the ground (Figure S2). Most branches below tree canopies had been pruned or died off naturally (Figure S2).

**Figure 2.**
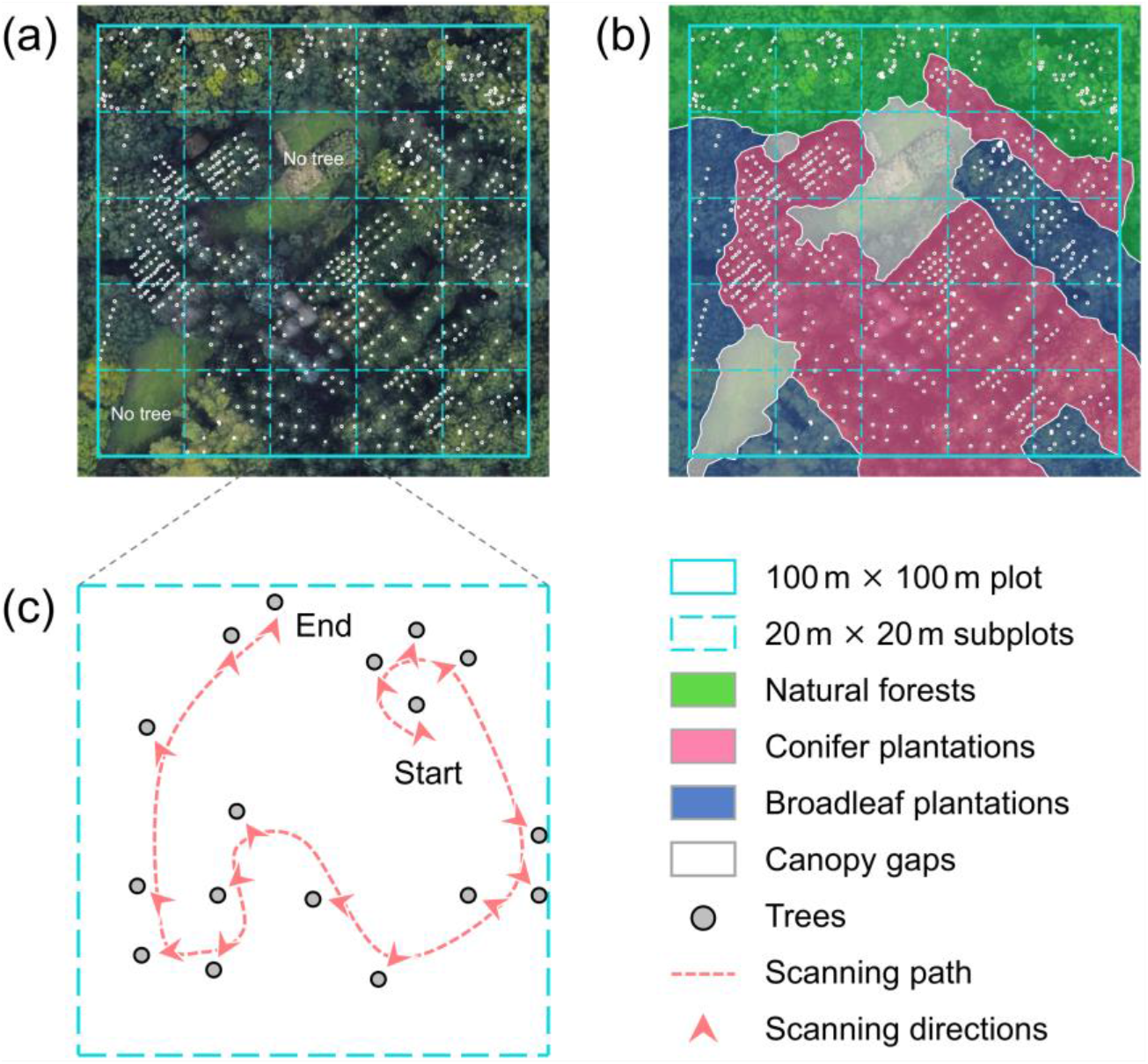
Overview of the study plot. (a, b) The 100 m × 100 m plot consisting of 20 m × 20 m subplots and multiple forest types. (c) An example of the path walked during scanning and the directions from which trees were scanned by ForestScanner. Background image: Google Earth.

All the trees with DBH ≥5 cm were measured and mapped using an iPhone 13 Pro and an iPad Pro 2021. We defined the 0.04 ha subplots as the unit of scanning, with consideration given to the plot sizes often adopted in national forest inventory programs (Paul, Kimberley, & Beets, 2019) and previous iPhone/iPad assessments which used 0.016 ha (Gollob et al., 2021) and 0.063 ha plots (Mokroš et al., 2021). Using each device, the surveyor scanned the trees consecutively in each subplot (Figure 2c). Each tree was scanned from one side. The geographic coordinates of the starting point were obtained by the GNSS receiver built into the iPhone/iPad. Scanning was conducted in 23 subplots since there were no trees in two subplots (Figure 2a).

To compare the results of ForestScanner with conventional methods, we measured the DBH using a diameter tape. The measurement was conducted by one measurer and one note-taker. We also measured the spatial coordinates of the trees using a combination of laser range-finder, electronic inclinometer, and electronic compass (Impulse 200 and MapStar Compass Module; Laser Technology Inc., Centennial, CO, USA) attached to a personal digital assistant (PA692; Unitech Electronics Co., Ltd., New Taipei City, Taiwan). Using this surveying system, two people conducted the survey; one person operating the system and the other holding a reflection board by the trees. We conducted closed traverses, starting from a control point (i.e., a corner of the plot, a midpoint of the plot edges, or the plot center) and either returning to the same point or closing on another control point. Trees were mapped by measuring the distances and angles from the traverse stations. The geographic coordinates of the control points were obtained by a multi-band GNSS mobile station D-RTK2 (SZ DJI Technology Co., Ltd., Shenzhen, China). The RTK-GNSS measurement at each point took ~5 seconds.

### 3.2 Data analyses

We compared the DBH measured by ForestScanner and diameter tape based on coefficients of determination (R^2^), root mean squared errors (RMSE), relative RMSE (%RMSE), biases, and relative biases (%biases). We also tested whether the amount of difference between DBH measured by the two methods changed with DBH. We used segmented regression to account for possible nonlinear relationships between DBH and the differences in DBH (Gollob et al., 2021)

We assessed the accuracy of tree mapping by ForestScanner based on the relative and absolute coordinates of trees. For relative coordinates, we compared the spatial distances among trees measured by ForestScanner and the conventional surveying system within each subplot. The comparisons were made using Mantel R^2^, a metric that quantifies the agreement between two distance matrices. For absolute coordinates, we calculated the distances between the coordinates of the same trees measured by ForestScanner and the conventional surveying system. Note that, while ForestScanner keeps track of the relative coordinates solely by IMU, the absolute coordinates are determined by both the built-in GNSS receiver and IMU. To assess the accuracy of the built-in GNSS receiver in determining the initial coordinates of each survey, we tested the relationships between the positioning errors of the first tree surveyed in each subplot and those of all trees.

### 3.3 DBH accuracy

ForestScanner successfully measured DBH of all the 672 trees (≥5 cm) in the plot. Tree detection rate was 100% since the trees were scanned independently on site. The DBH measured by ForestScanner and diameter tape showed good agreement; the R^2^ values were 0.963 for iPhone and 0.961 for iPad (Figure 3a, 3b). The RMSE and %RMSE were 2.3 cm and 10.3 % for iPhone and 2.3 cm and 10.5 % for iPad (Fig. 3c, 3d; Table S1). The biases and %biases were 0.2 cm and 0.8% for iPhone and 0.2 cm and 0.7% for iPad (Fig. 3c, 3d; Table S1). The DBH of small trees tended to be overestimated by iPhones/iPads (Fig. 3c, 3d), a pattern which was also observed in a previous study (Gollob et al., 2021). The DBH measured in conifer plantations, broadleaf plantations, and natural forests showed similar values for R^2^, RMSE, and biases (Table S1). The values were comparable with those reported in previous studies using iPhones/iPads: Detection rate = ~80%, R^2^ = 0.973, and RMSE = 3.1–4.5 cm (Gollob et al., 2021; Mokroš et al., 2021).

**Figure 3.**
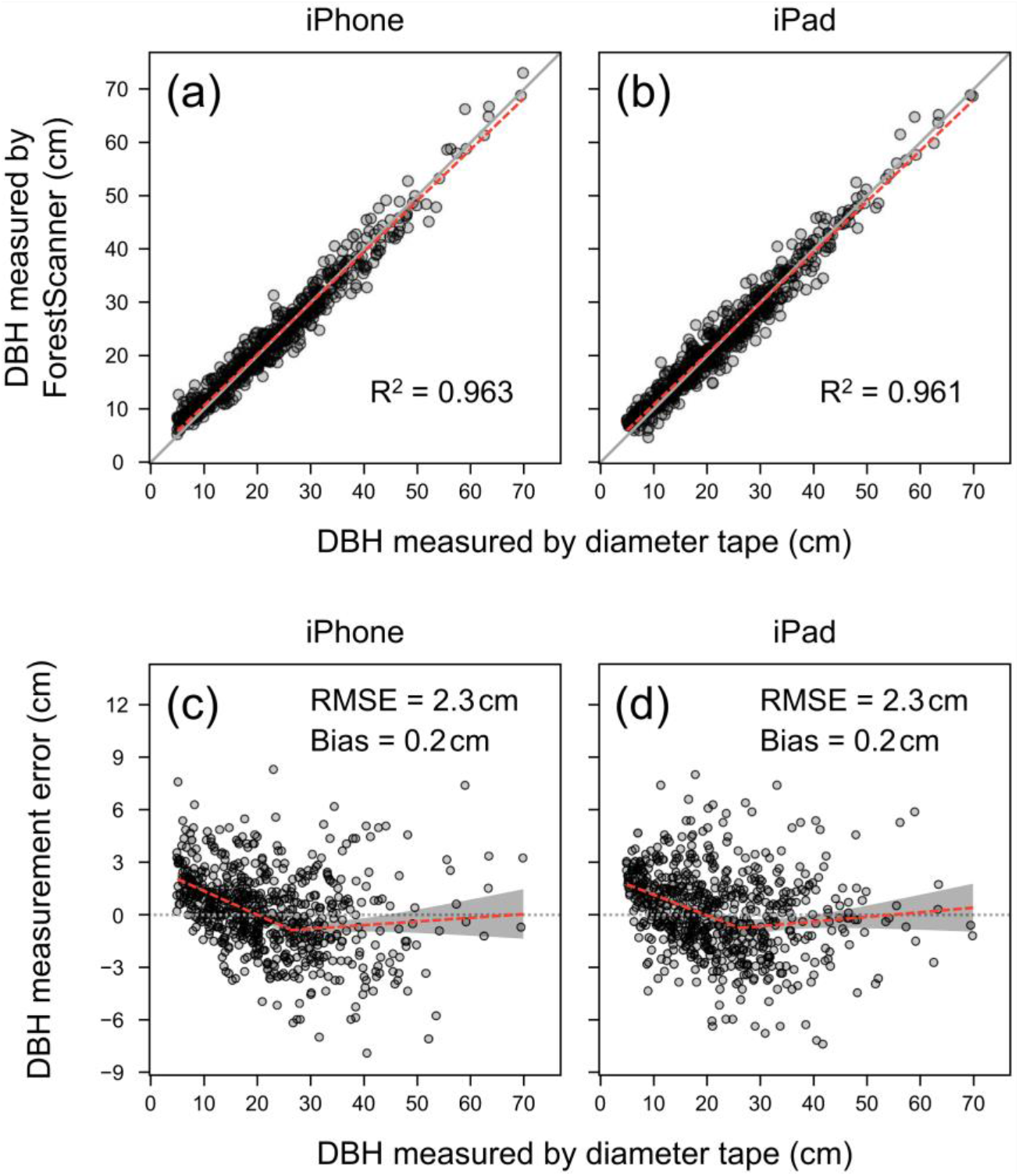
(a, b) Agreement between DBH measured by diameter tape (DBH_*x*_) and ForestScanner (DBH_*y*_). The red lines indicate linear regression models. (c, d) Relationships between DBHx and DBH measurement errors (DBH_*y*_ - DBH_*x*_). The red lines indicate segmented regression models.

### 3.4 Mapping accuracy

Spatial distances among trees measured by ForestScanner and the conventional surveying system were almost identical (Figure 4a, 4b). The Mantel R^2^ values, averaged across 23 subplots, were 0.999 for both iPhone and iPad (Figure 3c, 3d), indicating high accuracies of IMU in keeping track of the relative coordinates.

**Figure 4.**
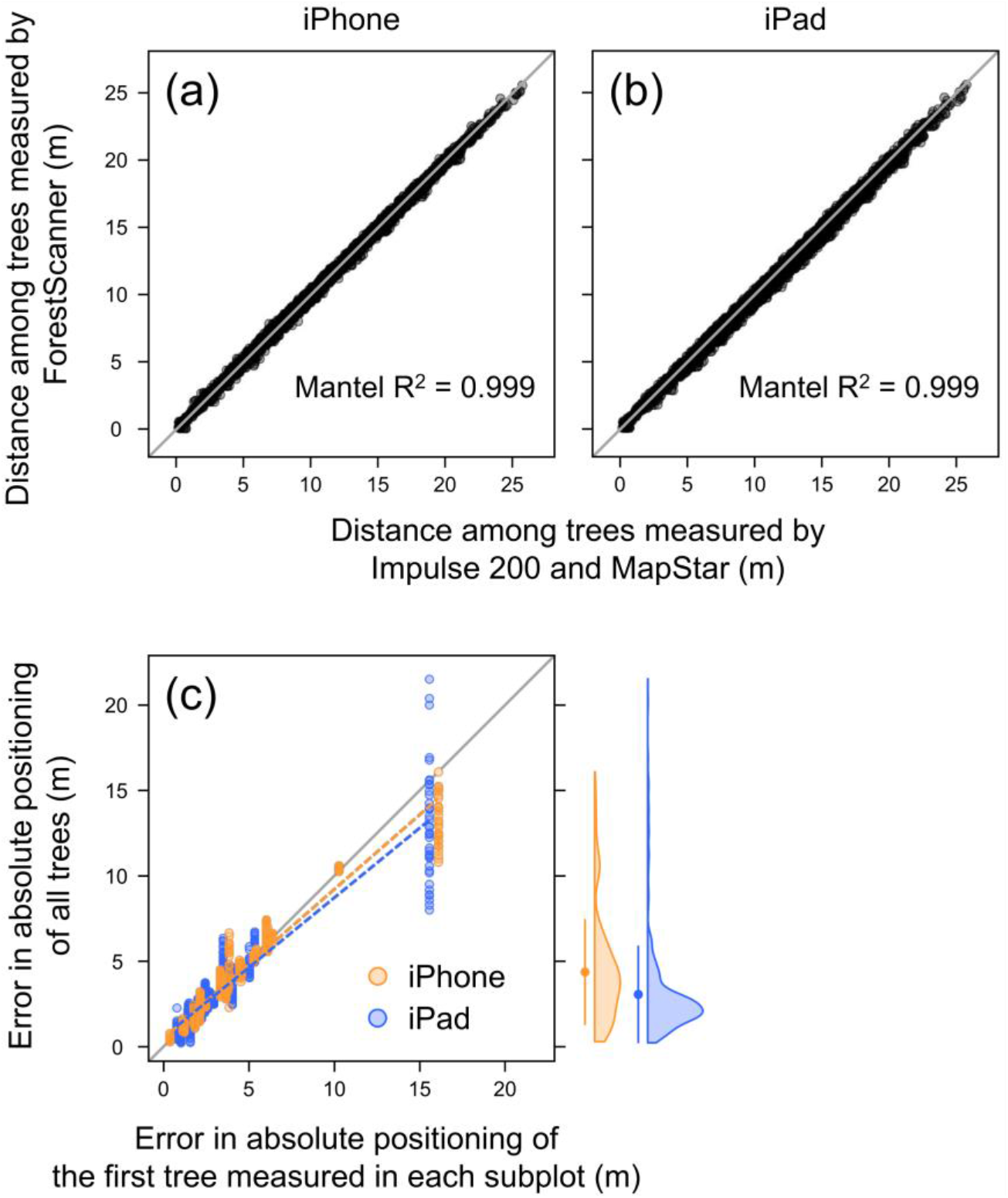
(a, b) Agreement between the distances among trees measured by a conventional surveying system (a combination of Impulse 200 and MapStar Compass Module) and ForestScanner. (c) Relationship between the absolute positioning errors of the first tree and all trees in each subplot. The dashed lines indicate linear regression models. The circles and bars on the right indicate mean ± SD.

The absolute positioning errors (i.e., distances between the coordinates of the same trees measured by ForestScanner and the conventional surveying system) were, on average, 4.4 m for iPhone and 3.1 m for iPad (Figure 4c). The absolute positioning errors were largely explained by the error of the first tree surveyed in each subplot (Figure 4c), indicating that the initial GNSS error caused shifts in the tree coordinates within each subplot (Figure S3). While we used the iPhone/iPad’s built-in GNSS receiver to determine the initial coordinates of each survey, the use of higher-accuracy GNSS receivers connectable to mobile devices (e.g., Trimble R1; Trimble Inc., Sunnyvale, CA, USA) could reduce the absolute positioning errors.

### 3.5 Time spent for scanning

Using an iPhone, it took 1 h 39 min (= 1.66 h) for one person to measure the DBH and coordinates of all the 672 trees in the plot (Table 1). It took 1 h 38 min (= 1.64 h) with an iPad to do the same things. Using a diameter tape, it took 3 h 13 min (= 3.21 h) for two people to measure the DBH (Table 1). To map the trees using the conventional surveying system, it took 8 h 53 min (= 8.88 h) for two people (Table 1). As such, compared to the conventional tools, ForestScanner reduced the person-hours (i.e., the number of workers × hours) required for measuring DBH to 25.7% (= 1.65/6.42), for mapping trees to 9.3% (= 1.65/17.76), and for doing both to 6.8% (= 1.65/24.18).

**Table 1.** The number of workers and time spent to measure the DBH and spatial coordinates of 672 trees in a 1 ha plot using ForestScanner (iPhone and iPad), a diameter tape, and a conventional surveying system (a combination of Impulse 200 and MapStar Compass Module).

|  | iPhone | iPad | Diameter tape | Impulse 200 and MapStar |
| --- | --- | --- | --- | --- |
| Items measured | DBH and coordinates | DBH and coordinates | DBH | Coordinates |
| Number of workers | 1 | 1 | 2* | 2† |
| Time spent (hours) | 1.66 | 1.64 | 3.21 | 8.88 |
| Workers × hours | 1.66 | 1.64 | 6.42 | 17.76 |

\* A measurer and a note-taker.
† An operator and a reflection-board holder.

## 4 DISCUSSION

ForestScanner is a mobile application for conducting forest inventories by means of LiDAR-equipped iPhones/iPads. We demonstrated that the application requires less labor and time in the field compared to using a dimeter tape and a conventional surveying method (Table 1). Tree diameters and coordinates measured by ForestScanner and the conventional tools showed good agreement (Figures 3 and 4). Moving forward, further research comparing ForestScanner with other tools are needed. For example, while we used a diameter tape in this study, modern tools such as computer calipers could be more comparable to ForestScanner in terms of time-persons required for measurements. In addition, as of yet, we have applied ForestScanner only in one site with a relatively low tree density and moderate or no understory vegetation (Figure S2). Future works should assess the performance of ForestScanner in various forest environments.

ForestScanner itself also has room for improvements. Specifically, the instance segmentation model (YOLACT++; Bolya et al., 2022) used in ForestScanner has, so far, been trained with images of tree stems without surrounding objects such as branches, lianas, or understory vegetation (Figure S1). Training the model to distinguish such objects from stems could reduce potential occlusions. Another challenge is that the scanning distance of iPhones/iPads is currently limited to ≤5 m. A promising way forward is to make some longer-distance hand-sized LiDAR units (e.g., Livox Mid-70; Livox Technology Co. Ltd., Shenzhen, China) attachable to iPhones/iPads. Such external LiDAR should allow ForestScanner to measure tree diameters from further distances and other forest metrics including tree heights.

In this study, we introduced ForestScanner, a free iPhone/iPad application that enables on-site, real-time measurements of tree diameters and spatial coordinates. The application is intuitively operable and does not require any manual post-analyses of 3D point clouds, providing access to LiDAR technology for non-experts including students and citizen scientists. Future research needs to validate the application in various forest types across different biomes. With further developments of internal models and add-on hardware (e.g., external LiDAR units), the utility of ForestScanner is expected to grow. We believe that ForestScanner will facilitate LiDAR-based resource assessments and biodiversity monitoring in forests worldwide.

## AUTHORS’ CONTRIBUTIONS

ST conceived the study. KY developed the application with inputs from ST and NF. ST and NF conducted the fieldwork. ST analyzed the data and wrote the manuscript. All authors contributed to manuscript revisions.

## DATA ACCESSIBILITY

Tree diameter and coordinate data obtained by iPhone, iPad, and conventional methods will be made available at FigShare upon acceptance. A demonstration of ForestScanner usage (Video S1) is available at https://doi.org/10.6084/m9.figshare.17161823.v1.

## ACKNOWLEDGEMENTS

We thank Atticus Stovall and two anonymous reviewers for constructive comments. This work was supported by the Japan Society for the Promotion of Science (Grant-in-Aid for Early-Career Scientists 21K14880) and the Council for the Development of Smart Forestry EZO Model.

## Notes

### Competing Interest Statement

The authors have declared no competing interest.

## REFERENCES

Bolya, D., Zhou, C., Xiao, F., & Lee, Y. J. (2022). YOLACT++: Better real-time instance segmentation. IEEE Transactions on Pattern Analysis and Machine Intelligence, 44, 1108–1121.

Bu, G., & Wang, P. (2016). Adaptive circle-ellipse fitting method for estimating tree diameter based on single terrestrial laser scanning. Journal of Applied Remote Sensing, 10, 026040.

Bunting, P., Armston, J., Lucas, R. M., & Clewley, D. (2013). Sorted pulse data (SPD) library. Part I: A generic file format for LiDAR data from pulsed laser systems in terrestrial environments. Computers and Geosciences, 56, 197–206.

Calders, K., Newnham, G., Burt, A., Murphy, S., Raumonen, P., Herold, M.,…Kaasalainen, M. (2015). Nondestructive estimates of above-ground biomass using terrestrial laser scanning. Methods in Ecology and Evolution, 6, 198–208.

Corona, P., Chirici, G., McRoberts, R. E., Winter, S., & Barbati, A. (2011). Contribution of large-scale forest inventories to biodiversity assessment and monitoring. Forest Ecology and Management, 262, 2061–2069.

Dassot, M., Constant, T., & Fournier, M. (2011). The use of terrestrial LiDAR technology in forest science: Application fields, benefits and challenges. Annals of Forest Science, 68, 959–974.

Eitel, J. U. H., Vierling, L. A., & Magney, T. S. (2013). A lightweight, low cost autonomously operating terrestrial laser scanner for quantifying and monitoring ecosystem structural dynamics. Agricultural and Forest Meteorology, 180, 86–96.

Gollob, C., Ritter, T., Kraßnitzer, R., Tockner, A., & Nothdurft, A. (2021). Measurement of forest inventory parameters with Apple iPad Pro and integrated LiDAR technology. Remote Sensing, 13, 3129.

Kunstler, G., Falster, D., Coomes, D. A., Hui, F., Kooyman, Robert, M., Laughlin, D. C.,…Ruiz-Benito, P. (2015). Plant functional traits have globally consistent effects on competition. Nature, 529, 1–15.

Liang, X., Kankare, V., Hyyppä, J., Wang, Y., Kukko, A., Haggrén, H.,…Vastaranta, M. (2016). Terrestrial laser scanning in forest inventories. ISPRS Journal of Photogrammetry and Remote Sensing, 115, 63–77.

Mokroš, M., Mikita, T., Singh, A., Tomaštík, J., Chudá, J., Wężyk, P.,…Liang, X. (2021). Novel low-cost mobile mapping systems for forest inventories as terrestrial laser scanning alternatives. International Journal of Applied Earth Observation and Geoinformation, 104, 102512.

Newnham, G. J., Armston, J. D., Calders, K., Disney, M. I., Lovell, J. L., Schaaf, C. B.,…Mark Danson, F. (2015). Terrestrial laser scanning for plot-scale forest measurement. Current Forestry Reports, 1, 239–251.

Paul, T. S. H., Kimberley, M. O., & Beets, P. N. (2019). Thinking outside the square: Evidence that plot shape and layout in forest inventories can bias estimates of stand metrics. Methods in Ecology and Evolution, 10, 381–388.

Pueschel, P., Newnham, G., Rock, G., Udelhoven, T., Werner, W., & Hill, J. (2013). The influence of scan mode and circle fitting on tree stem detection, stem diameter and volume extraction from terrestrial laser scans. ISPRS Journal of Photogrammetry and Remote Sensing, 77, 44–56.

Tatsumi, S., Cadotte, M. W., & Mori, A. S. (2019). Individual-based models of community assembly: Neighbourhood competition drives phylogenetic community structure. Journal of Ecology, 107, 735–746.

Tatsumi, S., Iritani, R., & Cadotte, M. W. (2021). Temporal changes in spatial variation: Partitioning the extinction and colonisation components of beta diversity. Ecology Letters, 24, 1063–1072.

Tatsumi, S., Owari, T., & Mori, A. S. (2016). Estimating competition coefficients in tree communities: A hierarchical Bayesian approach to neighborhood analysis. Ecosphere, 7, e01273.

Vogt, M., Rips, A., & Emmelmann, C. (2021). Comparison of iPad Pro®’s LiDAR and TrueDepth capabilities with an industrial 3D scanning solution. Technologies, 9, 25.

