## Supporting Information for "ForestScanner: A mobile application for measuring and mapping trees with LiDAR-equipped iPhone and iPad"

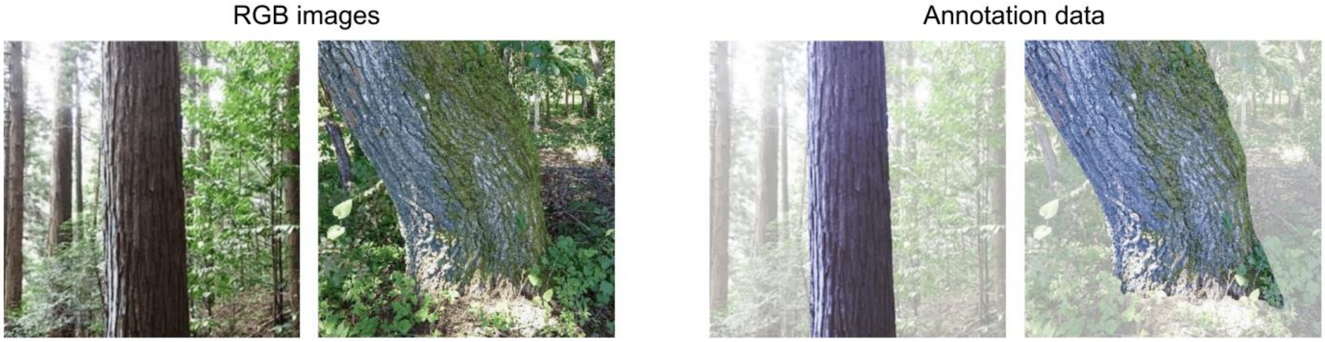


**Figure S1**. Examples of RGB images and annotation data used to train the instance segmentation model (YOLACT++) in ForestScanner.

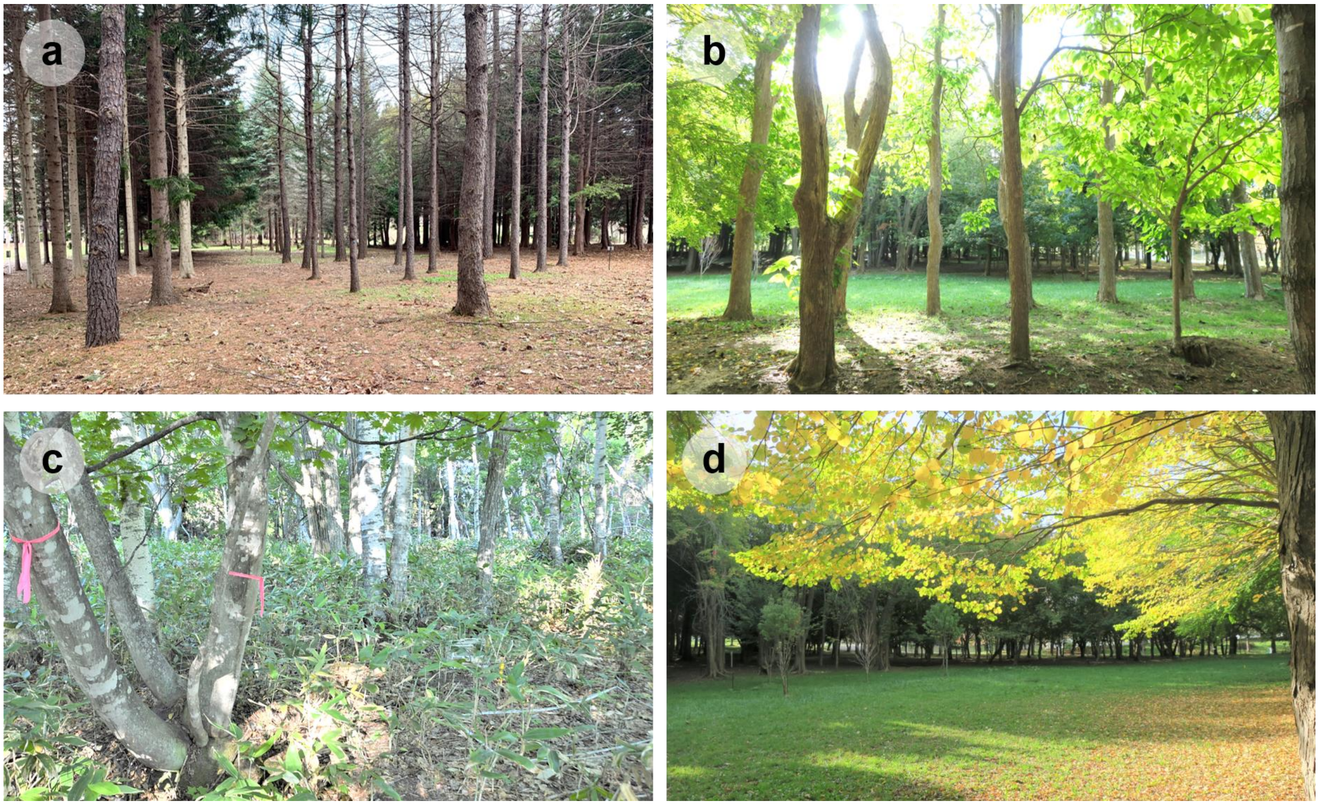


**Figure S2**. Forest types in the study plot: (a) Conifer plantation, (b) broadleaf plantation, (c) natural forest, and (d) forest gap. Pictures a, b, and d were taken in the study plot. Picture c was taken in a naturel forest ~1.7 km away from the plot and is presented here due to a lack of picture in the study plot. The natural forests in picture c and the study plot had the same stand age (regenerated after the same forest fire in 1912) and understory vegetation (dwarf bamboos: *Sasa senanensis*). In the plantations, branches had been pruned up to ~4 m in height from the ground. In natural forests, most of the branches below tree canopies had died off. Photo credits: (a) Tatsuya Sasaki, (b) and (d), Hirotada Kobayashi, and (c) Shinichi Tatsumi.

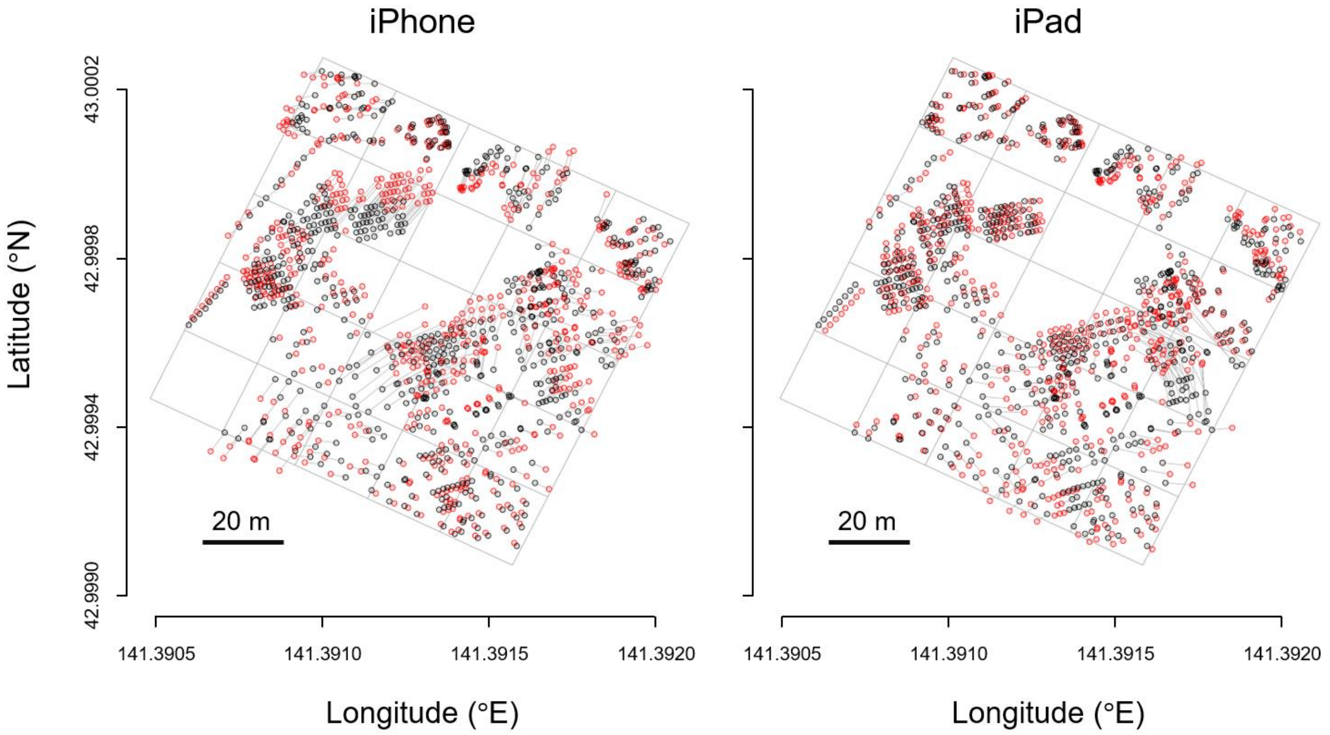


**Figure S3**. Spatial coordinates of trees measured by iPhone and iPad (red circles) vs. a conventional surveying system (black circles) in the 1 ha study site. Grey lines connecting the circles indicate the locations of the same tree measured by two different methods. The grids show 20 m × 20 m subplots.

**Table S1**. The $R^2$, RMSE, %RMSE, Bias, and %Bias between the DBH measured by iPhone and iPad vs. a diameter tape in three different forest types.

| Forest type | Number of trees (*n*) | iPhone | | | | | iPad | | | | |
|---|---|---|---|---|---|---|---|---|---|---|---|
| | | $R^2$ | RMSE (cm) | %RMSE (%) | Bias (cm) | %Bias (%) | $R^2$ | RMSE (cm) | %RMSE (%) | Bias (cm) | %Bias (%) |
| Conifer plantation | 411 | 0.959 | 2.1 | 8.8 | −0.3 | −1.2 | 0.947 | 2.4 | 10.0 | −0.1 | −0.6 |
| Broadleaf plantation | 118 | 0.971 | 2.4 | 11.5 | 0.4 | 2.1 | 0.979 | 2.0 | 9.7 | 0.2 | 1.0 |
| Natural forest | 143 | 0.970 | 2.4 | 12.6 | 0.4 | 2.3 | 0.973 | 2.0 | 10.6 | 0.2 | 1.1 |
| Total | 672 | 0.963 | 2.3 | 10.3 | 0.2 | 0.8 | 0.961 | 2.3 | 10.5 | 0.2 | 0.7 |
